# Interplay between canonical Wnt signaling and α5β1 integrins modulates mechanoresponse in human articular cartilage

**DOI:** 10.64898/2026.05.20.726448

**Authors:** N Viudes-Sarrión, R Castro-Viñuelas, N Vaes, EJ Blain, RJ Lories, I Jonkers

## Abstract

**Objectives:** Mechanical cues are essential for maintaining cartilage function, yet how they integrate with molecular pathways dysregulated in osteoarthritis (OA) remains poorly defined in human tissue. Canonical Wnt signalling influences cartilage biology and cell–matrix interactions, but its role in integrin-dependent mechanoregulation in human cartilage is not fully understood. This study aimed to determine how Wnt activation affects chondrocyte responses to physiological mechanical loading, with a focus on α5β1integrin and cytoskeletal organisation.

**Methods:** Human cartilage explants from non-OA and OA donors were subjected to short-term physiological cyclic compression. Canonical Wnt signalling was activated with CHIR99021, and integrin-mediated adhesion was modulated using the α5β1 blocking peptide ATN-161 during loading. Chondrocyte responses were assessed by analysing mechanoresponsive and matrix-related gene expression, α5β1 complex formation via proximity ligation assay and actin cytoskeletal organisation by confocal microscopy.

**Results:** OA chondrocytes exhibited a distinct integrin profile, characterised by increased *ITGA5* and *ITGB1* but reduced *ITGA10* expression. In non-OA cartilage, canonical Wnt activation increased *ITGB1* expression and α5β1 integrin complex formation, while mechanical loading further enhanced *ITGA5* and *ITGB1* transcription under Wnt-activated conditions. Under control conditions, loading induced mechanoresponsive and anabolic gene expression in non-OA cartilage; these responses were attenuated following Wnt-activation and partially restored by α5β1 blockade. Mechanical loading induced F-actin reorganization toward a more cortical distribution across cartilage zones, irrespective of disease status or treatment. Wnt activation did not result in distinct cytoskeletal phenotypes under load, and load-induced actin remodelling was comparable between groups.

**Conclusion:** These findings identify α5β1integrin as a key mediator linking canonical Wnt signalling to altered chondrocyte mechanoresponsiveness in human cartilage. While mechanical loading consistently induced cortical F-actin reorganization, Wnt-associated changes in load responsiveness arose primarily from integrin-dependent mechanisms rather than major alterations in actin organization. This study highlights the complexity of cartilage mechanoregulation and identifies integrin-mediated signaling as important contributors to canonical Wnt-driven alterations in load responsiveness relevant to OA.

## INTRODUCTION

Osteoarthritis (OA), a progressive joint disease characterized by articular cartilage damage, synovial inflammation and subchondral bone remodeling, one of the leading causes of disability worldwide arising from a multifactorial interplay of mechanical, biochemical, and genetic factors^1^. A hallmark of OA pathophysiology is an altered cartilage response to mechanical loading^2^. Under physiological conditions, mechanical stimuli are essential for maintaining cartilage homeostasis^2^, whereas in OA this mechanoresponsive behavior becomes dysregulated, favoring catabolic signaling pathways and tissue degradation^3,4^.

Chondrocytes sense and respond to mechanical forces through mechanotransduction pathways, with integrins as key mediators. These heterodimeric receptors connect the extracellular matrix (ECM) to the actin cytoskeleton and facilitate bidirectional signaling essential for load-dependent matrix maintenance^5,6^. Several integrins, including α5β1 or αvβ3, regulate chondrocyte–matrix interactions, with α5β1 identified as a principal chondrocyte mechanoreceptor^5^. In healthy cartilage, integrin-mediated signaling coordinates cytoskeletal tension and anabolic activity^5^, whereas in OA altered integrin expression and activation have been associated with aberrant downstream signaling, resulting in enhanced mitogen-activated protein kinase (MAPK) and nuclear factor κB (NF-κB) signaling, increased matrix metalloproteinase (MMP) production and ECM breakdown^7,8^.

Canonical Wnt signaling, a key regulator of cell differentiation and tissue homeostasis, is aberrantly activated in OA^9,10^. Excessive Wnt activity induces catabolic gene programs, chondrocyte hypertrophy, and accelerates matrix degradation^11^. Consistently, we previously showed that canonical Wnt activation alters the chondrocyte mechanoresponse in human cartilage^12^. Dysregulated Wnt signaling also influences mechanobiological responses in other tissues and cell types, including bone and fibroblasts, as well as in tumor progression, where Wnt and integrin pathways converge at shared signaling hubs, such as the FAK–Src axis and β-catenin stabilization^13,14^

Although both integrin and Wnt pathways are dysregulated in OA, it remains unclear whether altered Wnt activity directly disrupts cartilage mechanotransduction through effects on the integrin– cytoskeleton axis. Specifically, the extent to which canonical Wnt signaling modifies integrin-mediated force transmission in human cartilage has not been defined. We hypothesized that impaired mechanosensitivity in OA cartilage may arise from maladaptive interactions between canonical Wnt signaling and the integrin-cytoskeleton axis. To test this, we experimentally activated canonical Wnt signaling in healthy human cartilage explants, to induce molecular features associated with an OA-like phenotype and examined how this modulation affects chondrocyte integrin expression and responses to physiological loading. As cytoskeletal remodelling is essential for force transmission, we further examined whether Wnt-induced alterations in mechanoresponsiveness were associated with changes in F-actin organization. Based on previous work showing that physiological loading promotes cortical actin distribution, whereas impaired mechanotransduction disrupts this arrangement^15,16^, we examined whether Wnt-induced alterations in mechanoresponsiveness were associated with changes in F-actin organisation.

## METHODS

### Human articular cartilage explants and primary chondrocyte collection and culture

Human articular cartilage samples were obtained from patients undergoing hip replacement surgery, as approved by the UZ Leuven Ethics and Biobank Committee (S56271). Cartilage tissue was rinsed in phosphate-buffered saline (PBS) supplemented with 1% antibiotic/antimycotic (Gibco), and 8 mm-diameter explants were generated using a disposable biopsy punch (Robbins Instruments). Explants were cultured for 24h at 37°C and 5% CO_2_ in DMEM/F12 (Gibco) supplemented with 10% fetal bovine serum (Gibco), 2 mM L-glutamine (Gibco), and 100units/mL penicillin, 100µg/mL streptomycin, and 0.25µg/mL amphotericin B (Gibco). For isolation of primary human articular chondrocytes (hACs), cartilage was dissected, rinsed, minced, and digested with pronase (2mg/mL Roche) for 90 min, followed by overnight digestion with collagenase B (1.5mg/mL; Roche). The resulting cell suspension was filtered through a 70µM strainer, and cells expanded under standard culture conditions in the same medium described above. Cartilage explants were obtained from 26 non-OA donors, undergoing hip replacement surgery for osteoporotic or malignancy-associated fracture, and from 6 patients with established OA. Sample size was defined by human tissue availability; however, experimental designs were paired within donors, and statistical models accounted for inter⍰donor variability through inclusion of donor as a random effect. Donor allocation, including sex and age distribution for each analysis, is detailed in Supplementary Table 1.

### Wnt activation and α5β1 integrin blocking experiments

Articular cartilage explants from seven non-OA donors were treated with Wnt agonist CHIR99021 (6µM; Sigma-Aldrich) or its vehicle (1µL/mL dimethyl sulfoxide, DMSO-control group) for 24h prior to mechanical stimulation. CHIR99021 concentration was selected based on previous reports^17^ and our previous work^12^. To block α5β1 integrin, the non-RGD-based integrin-binding peptide ATN-161 (Selleck Biotechnology) was used. A dose–response curve (100, 150 and 200nM; 4h incubation) was performed in non-OA explants based on concentrations reported in the literature^18^. Effective integrin blocking was validated by assessing tissue mechanoresponsiveness through *c-JUN, cFOS* and *SOX9* transcription (Supplementary Figure 2); the lowest effective concentration (100nM) was selected for subsequent experiments.

### Cartilage explant viability

As cartilage explant viability following CHIR99021 treatment was previously validated^12^, viability was assessed only following ATN-161 treatment. Three non-OA explants were treated with the highest ATN-161 concentration tested (200nM for 4h) and viability assessed by incubating explants with 10% PrestoBlue™ Cell Viability Reagent (Thermo Fisher) in culture medium for 2 h at 37°C (Supplementary Figure 1). Absorbance was measured at 570 nm and 600 nm using a BioTek Synergy™ HT microplate reader.

### Mechanical stimulation

Physiological mechanical stimulation was applied using a displacement-controlled compression protocol as previously described^12^. Briefly, cartilage explants were subjected to cyclic compression (10% strain, 1Hz)^19,20^ in two loading cycles of 1h separated by 1h of free swelling using an ElectroForce® BioDynamic 5210 bioreactor (TA Instruments) at 37°C and 5% CO_2_. Unloaded controls were maintained under identical free-swelling conditions. Mechanical loading was applied in three different experiments: (i) analysis of integrin expression during loading with and without Wnt activation; (ii) evaluation of whether α5β1 integrin blockade modifies Wnt-induced alterations in mechanoresponsiveness; and (iii) assessment of actin cytoskeletal reorganization immediately and 1h after loading under Wnt activation and/or α5β1 integrin blockade.

### RNA extraction

Following loading, cartilage explants were washed, snap-frozen and RNA isolated as previously described^12^. Briefly, frozen tissue was mechanically homogenized in TRIzol™ (Invitrogen) using CKMix50-R tubes in a Precellys Evolution homogenizer, followed by phase separation and RNA isopropanol/LPA precipitation with centrifugation. For primary hACs, lysis was performed directly in TRIzol™ prior to RNA extraction as described. RNA concentration and purity were assessed using NanoDrop spectrophotometry (ThermoFisher).

### Gene expression analysis

cDNA was synthesized using the RevertAid H Minus First Strand cDNA Synthesis Kit (Thermo Fisher) according to the manufacturer’s instructions. Quantitative PCR was performed using a StepOne Real-Time PCR System with Maxima SYBR Green qPCR Master Mix (Thermo Fisher). Relative gene expression was calculated using the comparative Ct method and normalized to the housekeeping gene *29S*. PCR conditions consisted of 40 amplification cycles of 15s at 95°C and 45s of 60°C. Melting curve analysis was performed to confirm amplification specificity. Primer sequences are listed in Supplementary Table 2.

### Proximity Ligation Assay (PLA)

Non-OA and OA primary hACs were seeded at passage 0 (P0) on fibronectin-coated coverslips (2 µg/cm^2^; Sigma-Aldrich). Non-OA cells were treated with CHIR99021 or vehicle prior to fixation, OA cells were analyzed under basal conditions. After one week in culture, α5β1 integrin complex formation was assessed using the Duolink® *In Situ* Red Starter Kit (Sigma-Aldrich) following manufacturer recommendations. Briefly, cells were fixed with 4% paraformaldehyde (10min) and permeabilized with 0.1% Triton X-100 (10min). After blocking for 60 min at 37°C, cells were incubated overnight at 4°C with primary antibodies against α5 integrin (rabbit monoclonal, clone EPR7854; Abcam, ab150361) and β1 integrin (mouse monoclonal, clone P5D2; Santa Cruz Biotechnology, sc-70809), each diluted 1:50. PLA probes (anti-rabbit MINUS and anti-mouse PLUS) were applied for 60 min at 37°C, followed by ligation (30min) and amplification (60min). Negative controls were performed by omitting primary antibodies while including PLA probes, ligation, and amplification steps. After washing, cell membranes were labelled using CellMarker Plasma Membrane Stain (Thermo Fisher) and coverslips mounted with VECTASHIELD® HardSet™ with DAPI.

Confocal images were acquired using a Nikon C2 microscope with a 40× objective. Z-stacks were collected at 0.3µm intervals using identical acquisition settings (405nm for DAPI, 561nm for PLA, 488nm for cell marker). Five independent fields were imaged per condition.

### Quantification of PLA

PLA images were analysed using a custom MATLAB-based script (https://gitlab.kuleuven.be/MAtrix/nele/pla_quantification). Nuclei were segmented from the DAPI channel, cell boundaries identified using the membrane marker, and PLA puncta detected in the red channel. Automatic segmentation was reviewed for each cell, with manual correction performed when necessary. Minimum puncta size threshold was fixed across all samples. Operators confirmed inclusion or exclusion of individual cells to ensure continuous quality control.

### Analysis of F-actin organisation

Following mechanical stimulation, explants were fixed overnight at 4°C in 4% paraformaldehyde, cryoprotected by sequential sucrose incubations (25–100%), embedded in OCT (Tissue-Tek), snap-frozen, and sectioned at 20µm thickness. Sections were mounted on Superfrost Plus™ slides (Thermo Fisher) and stored at −80°C. Sections were permeabilized in 0.5% Triton X-100 for 10 min, blocked in 5% BSA for 30 min, and stained with Alexa Fluor™ 635–conjugated phalloidin (1:400; Invitrogen, A34054) for 1h at room temperature. Sections were mounted with VECTASHIELD® HardSet™ with DAPI.

Confocal imaging was performed using a Zeiss LSM880 Airyscan microscope with a 63× oil-immersion objective (NA 1.4). Z-stacks were acquired at 0.35–0.50µm intervals using sequential excitation (405nm for DAPI, 633nm for phalloidin). Images were collected from three representative regions per cartilage zone (superficial and deep) per experimental condition.

### Quantification of F-actin distribution

F-actin distribution was quantified using a custom MATLAB script (https://gitlab.kuleuven.be/MAtrix/nele/actin_quantification). Individual cells were segmented and manually reviewed. The cortical region was defined as the outermost 10% of the cell area^21^, and the ratio of cortical to total F-actin intensity calculated by dividing the sum of pixel intensities within the cortical region by the total cell area, enabling normalized comparisons across cells of varying sizes.

### Statistics

Statistical analyses were performed with GraphPad Prism version 8 and R-Studio Version 4.5.1 with relevant packages.

To compare basal mRNA expression levels of selected integrin subunits between primary OA and non-OA hACs, independent two-sample, two-tailed t-tests with Welch’s correction were performed. Results are reported as differences of the means with corresponding 95% confidence intervals and P values.

For all other experiments: (i) integrin gene expression during mechanical loading and Wnt activation;, (ii) cartilage mechanoresponsive gene expression upon Wnt activation and/or integrin blocking, (iii) α5β1 integrin complex quantification, and (iv) cytoskeletal organization evaluation, data were analysed using linear mixed-effects models (LMM) (Supplementary Table 4) to account for the hierarchical structure of the data. Estimated marginal means (emmeans) were calculated, and pairwise comparisons were performed to evaluate differences between conditions. For experiments with multiple conditions, model-derived main effects and interactions were assessed using analysis of variance (ANOVA) on the fixed effects. Where appropriate, Tukey adjustment was applied for multiple comparisons within the model.

For (iii) and (iv) experiments, individual cells were treated as observations, and LMM were used to account for the non-independence of cells derived from the same donor, with donor included as a random effect. For (iii) experiments, pairwise comparisons were defined *a priori* based on the experimental design. As these comparisons were not exhaustive across all possible condition combinations, no correction for multiple testing was applied. For (iv) experiments, data were analysed separately for superficial and deep cartilage zones.

Results are reported as differences of the means with corresponding 95% confidence intervals and P-values. A P-value <0.05 was considered statistically significant. Effect sizes are reported on the log⍰transformed scale used for statistical analysis. All main effects and *post hoc* pairwise comparisons are reported in Supplementary Table 3.

## RESULTS

### Differential expression of selected integrin subunits in OA and non-OA human articular chondrocytes

Basal mRNA levels of selected integrin subunits were compared in primary OA and non-OA hAC at P0 (Figure 1). OA chondrocytes expressed significantly higher *ITGA5* (mean difference = 0.649, 95% CI [0.087–1.210], P = 0.0327) and *ITGB1* (mean difference = 0.340, 95% CI [0.014–0.666], P = 0.0445), encoding the fibronectin-binding α5β1 integrin. In contrast, expression of *ITGA10*, encoding the collagen type II–binding α10 subunit, was significantly reduced in OA chondrocytes (mean difference = −0.308, 95% CI [−0.510–−0.106], P = 0.0141). Expression levels of other integrin subunits did not differ significantly between OA and non-OA groups (Supplementary Table 3.1). Together, these findings indicate a selective shift in integrin subunit expression in OA chondrocytes, which may reflect alterations in cell–matrix interactions. Given the limited donor number, these analyses are exploratory and intended to identify candidate integrin subunits for subsequent mechanistic experiments.

**Figure 1.**
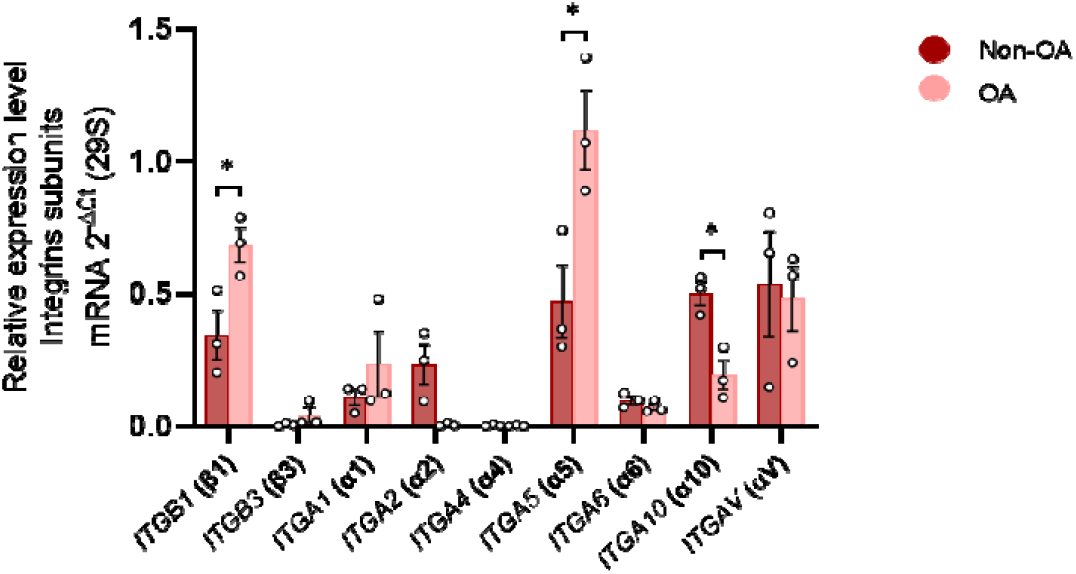
Differential expression of integrin subunits in human OA vs non-OA chondrocytes: Basal mRNA expression levels of integrin subunits were quantified in primary chondrocytes isolated from OA and non-OA donors. Data are presented as individual data points for each donor sample, along with the mean ± SEM (n = 3 per group); statistical significance was assessed using independent two-sample t-tests, two-tailed (P < 0.05 ).

### Wnt signaling activation modulates integrin expression in human cartilage tissue

To investigate whether canonical Wnt activation modulates integrin expression under basal and mechanically stimulated conditions, non-OA cartilage explants were treated with Wnt agonist CHIR99021 (CHIR) or vehicle (DMSO) and maintained under free-swelling or loading (Figure 2a). All statistical main effects and interaction terms derived from the LMM, as well as all *post hoc* pairwise comparisons are reported (Supplementary Table□3.2).

**Figure 2.**
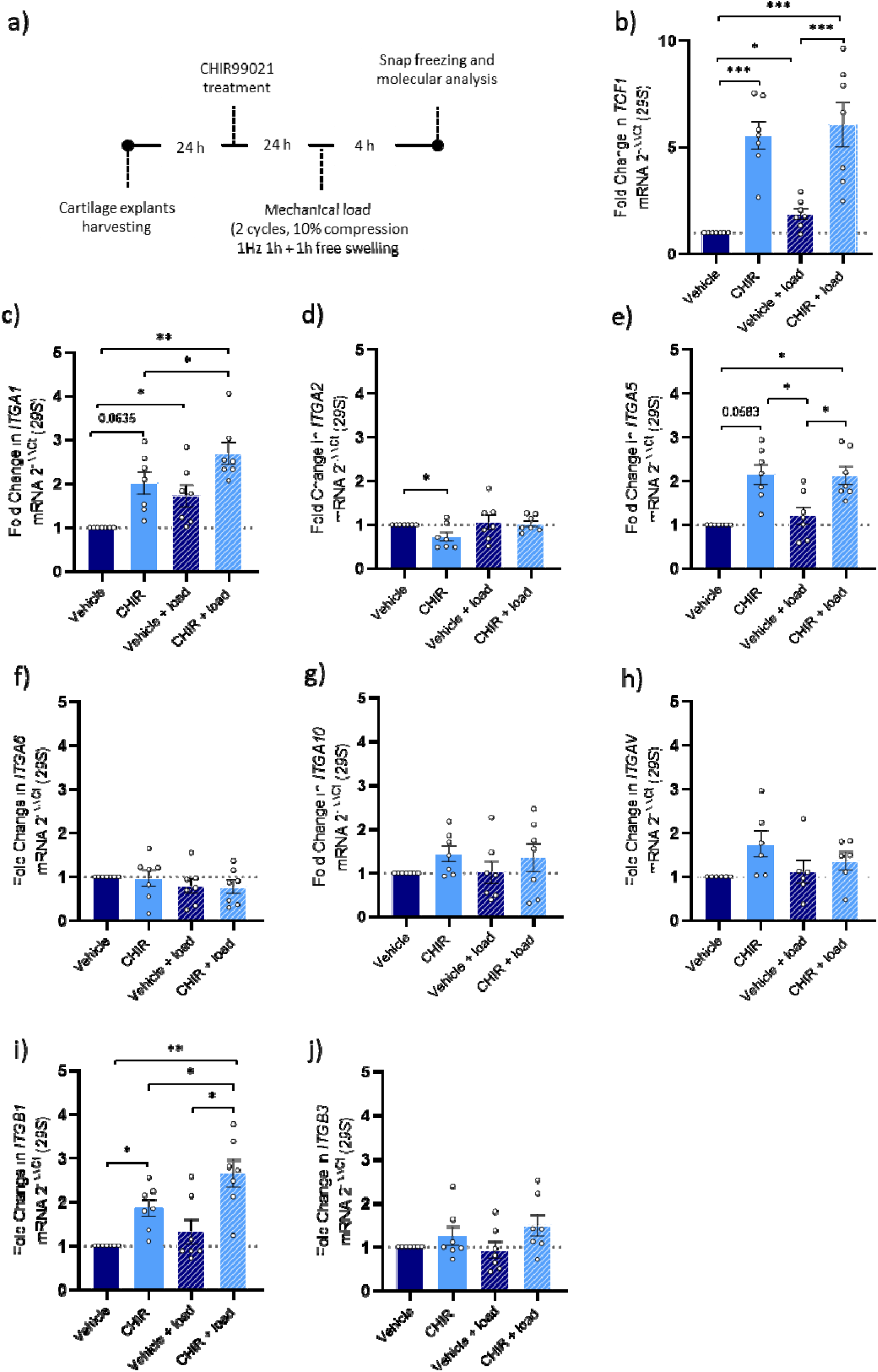
a) Schematic representation of the followed workflow. b) Real-time PCR analysis positive control gene *TCF-1* for canonical Wnt signaling activation, c-j) Real-time PCR analysis of integrin genes expressed in cartilage, including *ITGA1, ITGA5, ITGA10, ITGA2, ITGA6, ITGAV, ITGB1*, and *ITGB3* in human non-OA articular cartilage explants after mechanical stimulation (load) and/or treatment with 6μM CHIR99021 (CHIR) or vehicle. Data are presented as individual data points for each donor sample, along with the mean ± SEM for all treatment conditions (n = 7, P < 0.05, Tukey corrected tests in LMM). Vehicle = DMSO treatment, CHIR= CHIR99021 treatment, Load = mechanical loading.

Canonical Wnt activation was confirmed by increased expression of the Wnt target gene *TCF1* following CHIR99021 treatment (Figure 2b; *post hoc* pairwise comparison, mean difference = 0.723, 95% CI [0.501–0.946], P < 0.0001). Loading alone also induced *TCF1* expression relative to unloaded controls (mean difference = 0.249, 95% CI [0.026–0.471], P = 0.0245). When loading was applied in the presence of CHIR99021, *TCF1* levels were further increased compared with loading alone (mean difference = 0.488, 95% CI [0.266–0.711], P < 0.0001) (Figure 2b), indicating that mechanical stimulation can potentiate Wnt pathway activity.

Loading alone also significantly increased *ITGA1* expression compared with unloaded controls (mean difference = 0.235, 95% CI [0.012–0.471], P = 0.0374). CHIR99021 treatment significantly increased *ITGB1* expression compared with unloaded controls (mean difference = 0.180, 95% CI [−0.428– 0.670], P = 0.0400), and showed trends towards increased expression of *ITGA5* (mean difference = 0.229, 95% CI [−0.024–0.483], P = 0.0583) and *ITGA1* (mean difference = 0.149, 95% CI [−0.005– 0.373], P = 0.0635). Conversely, *ITGA2* expression was significantly reduced following CHIR99021 treatment (mean difference = −0.189, 95% CI [−0.373–−0.005], P = 0.0434) compared with unloaded controls.

When loading was applied under conditions of Wnt activation, *ITGA5* and *ITGB1* expression levels were significantly higher compared with loading alone (mean difference = 0.301, 95% CI [0.048– 0.555], P = 0.0171 and mean difference = 0.238, 95% CI [0.010–0.486], P = 0.0412, respectively) (Figure 2c). Furthermore, both *ITGA1* and *ITGB1* expression were significantly higher in the combined condition of CHIR99021 and loading compared with CHIR99021 alone (mean difference = 0.263, 95% CI [0.040–0.487], P = 0.0184 and mean difference = 0.232, 95% CI [0.016–0.479], P = 0.0404, respectively). The other integrin subunits were unaffected (Figure 2c).

These results show that in human cartilage, canonical Wnt activation modulates *ITGB1* and *ITGA2* and that loading amplifies these effects for *ITGA5, ITGB1* and *ITGA1* integrin expression, suggesting potential interactions between Wnt and integrins in mediating cartilage mechanoresponses.

### Wnt activation promotes α5β1 integrin complex formation in human chondrocytes

Given the observed upregulation of *ITGA5* and *ITGB1* in OA chondrocytes, as well as following Wnt stimulation in cartilage explants, we next examined whether these gene-level changes were associated with increased α5β1 heterodimer formation. α5β1 complexes were quantified by PLA in (i) primary OA hAC and (ii) non-OA hAC following CHIR99021 treatment. Main effects and a complete set of LMM pairwise comparisons is provided (Supplementary Table 3.3).

*Post hoc* pairwise comparisons revealed significantly higher α5β1 complex formation in OA compared with non-OA hAC (mean difference = 30.1, 95% CI [20.36–39.84], P = 0.0008) (Figure 3h). Similarly to OA conditions, α5β1 complex formation was increased in CHIR99021 treated non-OA hAC compared with vehicle controls (mean difference = 20.3, 95% CI [13.65–26.95], P < 0.0001) (Figure 3i).

**Figure 3.**
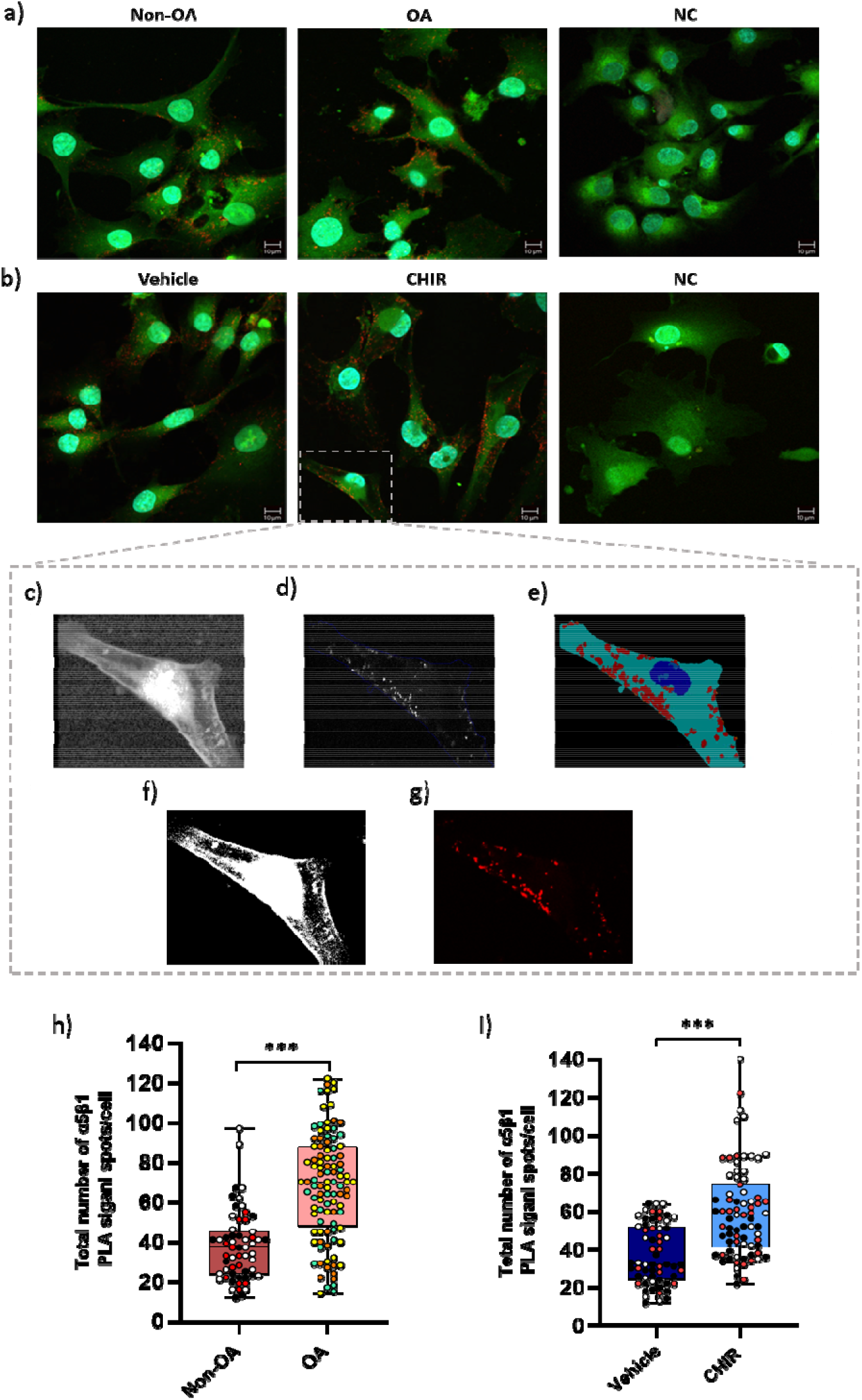
Increased formation of α5β1 integrin complexes in OA and Wnt-activated human chondrocytes (a) Representative images of the α5β1 PLA in hAC from non-OA and OA donors and a technical negative control used to confirm assay specificity. (b) Representative α5β1 PLA images from non-OA hAC treated with vehicle (control), non-OA hAC treated with 3 µM CHIR99021 to activate Wnt signaling, and their corresponding negative control. Nuclei are stained with DAPI (blue), Cell tracker labels the cytoplasm (green), and PLA puncta indicating α5β1 complex formation are shown in red. (c–e) Overview of the MATLAB-based analysis workflow used to quantify PLA signals: (c) the original cell tracker image used for cell identification; (d) PLA channel overlaid with automatically detected cell boundaries; (e) segmentation output showing cell masks (cyan), nuclear masks (blue), and PLA puncta segmentation (red). (f–g) ImageJ output used to validate segmentation and ensure correct interpretation of PLA puncta detection: (f) ImageJ cell tracker image; (g) ImageJ PLA channel. (h, i) Quantification of α5β1 PLA puncta per cell across two independent experimental comparisons: (h) non-OA versus OA donors, and (i) vehicle-treated versus CHIR99021-treated non-OA cells. Cellular PLA measurements were analysed using LLM-effects to account for the non-independence of cells derived from the same donor, with donor included as a random effect. Data are therefore interpreted at the donor level, and individual donors are represented by distinct colours in the dot plots (n=3). Given the use of predefined pairwise comparisons, no correction for multiple testing was applied. Scale bar: 10 µm. Vehicle = DMSO treatment, CHIR= CHIR99021 treatment.

These findings demonstrate that canonical Wnt activation increases α5β1 heterodimer formation in non-OA chondrocytes closely mirroring the increased abundance of α5β1 heterodimers observed in OA hAC.

### α5β1 integrin blockade partially mitigates Wnt-induced alterations in chondrocyte mechanoresponl**seness**

To determine whether CHIR99021-induced α5β1 increase contributes to impaired mechanoresponses, non-OA cartilage explants were treated with CHIR99021 and subjected to α5β1 blockade (ATN-161) prior to loading (Figure 4). Main effects and a complete set of LMM pairwise comparisons is provided (Supplementary Table 3.4).

**Figure 4.**
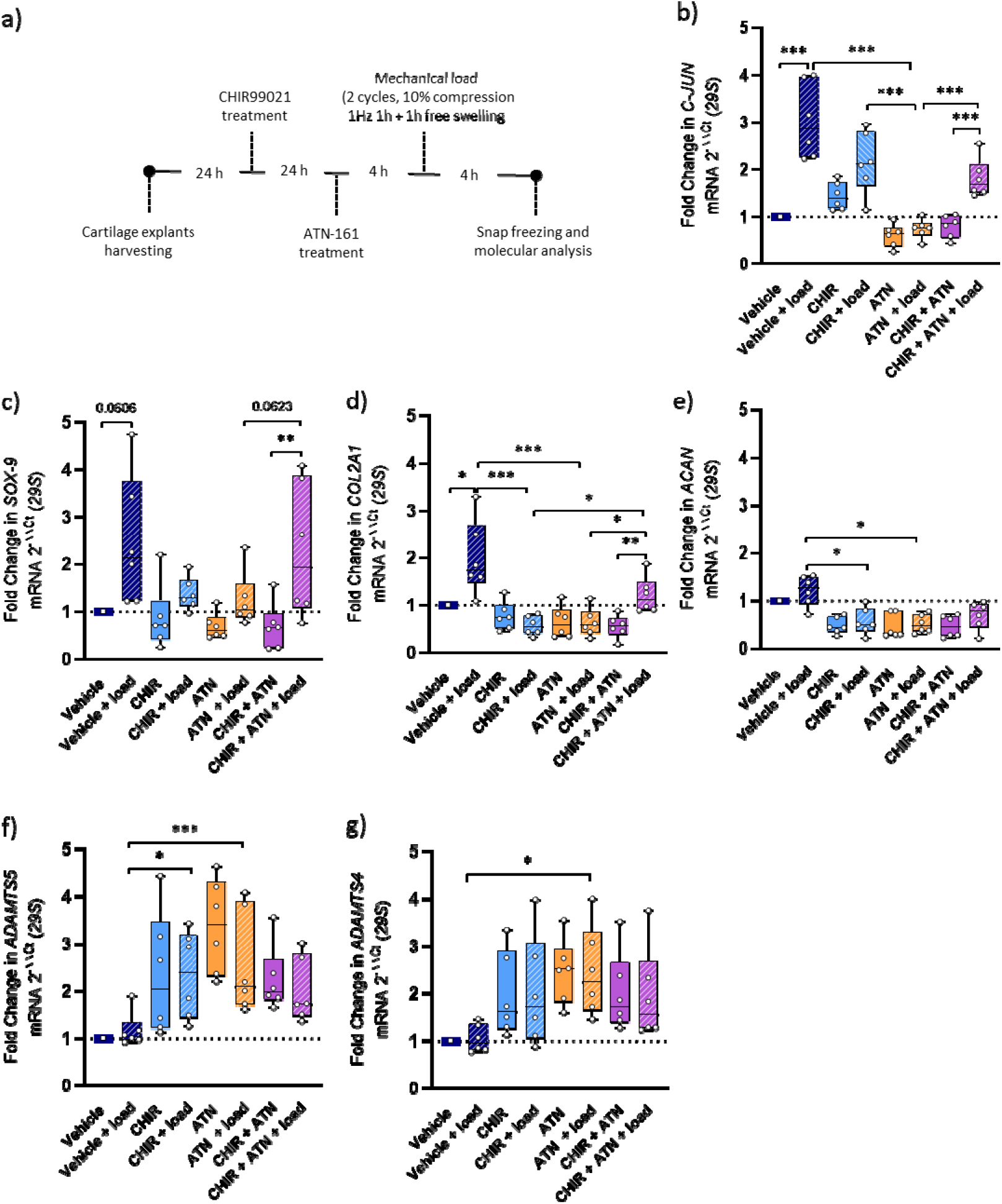
(a) Schematic representation of the experimental workflow, including unloaded and mechanically loaded explants treated with vehicle (DMSO), CHIR99021 (CHIR, 6 μM), the α5β1 integrin inhibitor ATN-161 (ATN, 100 μM), or their combinations. (b) Real-time PCR analysis of the mechanosensitive gene *c-JUN* across all unloaded and loaded conditions. (c–e) Real-time PCR analysis of the chondrogenic markers *SOX9, COL2A1*, and *ACAN* across all conditions. (f–g) Real-time PCR analysis of the catabolic markers *ADAMTS5* and *ADAMTS4* across all conditions. Data are presented as individual data points for each donor sample, along with the mean ± SEM for all treatment conditions. (n = 6, P < 0.05, Tukey corrected tests in LMM). Vehicle = DMSO; CHIR = CHIR99021; ATN = ATN-161; Load = mechanical loading.

As expected, *post hoc* pairwise comparisons revealed that loading alone increased transcription of mechanosensitive *c-JUN* (mean difference = 0.46, 95% CI [0.25–0.67], P < 0.0001) and the anabolic marker *COL2A1* (mean difference = 0.27, 95% CI [−0.04–0.58], P = 0.0132) compared to unloaded controls, with a trend towards increased *SOX-9* expression (mean difference = 0.33, 95% CI [−0.08– 0.74], P = 0.0606) (Figure 4b,c,d).

In contrast, Wnt activation during loading impaired this anabolic response, with decreased *COL2A1* (mean difference = −0.54, 95% CI [−0.85–−0.23], P < 0.0001) and *ACAN* expression (mean difference = −0.41, 95% CI [−0.74–−0.07], P = 0.0071) compared to loading alone (Figure 4d,e), and an increase of the catabolic marker *ADAMTS5* (mean difference = 0.26, 95% CI [0.01–0.52], P = 0.0471) (Figure 4f). CHIR99021 alone did not significantly alter these markers.

Blocking α5β1 during loading reduced transcription of *c-JUN* (mean difference = −0.61, 95% CI [−0.82–−0.40], P < 0.0001), *COL2A1* (mean difference = −0.51, 95% CI [−0.82–−0.20], P = 0.0001), and *ACAN* (mean difference = −0.45, 95% CI [−0.78–−0.11], P = 0.0022) compared with loading alone (Figure 4b,d,e). Furthermore, α5β1 blockade with loading (ATN + load) increased expression of the catabolic markers ADAMTS4 (mean difference = 0.37, 95% CI [0.10–0.64], P = 0.0020) and *ADAMTS5* (mean difference = 0.33, 95% CI [0.19–0.72], P = 0.001) compared to loading alone (Figure 4f,g).

Blocking α5β1 in CHIR99021-treated cartilage subjected to loading mitigated some of the Wnt-induced alterations observed. Compared with just blocking α5β1 and loading, *c-JUN* expression was significantly increased (mean difference = 0.39, 95% CI [0.18–0.60], P < 0.0001) and *SOX-9* showed a similar trend (mean difference = 0.21, 95% CI [−0.20–0.56], P = 0.0623) (Figure 4b,c). Importantly, blocking α5β1 integrins with ATN-161 in CHIR-treated explants subjected to loading significantly increased *COL2A1* expression compared to CHIR-treated loaded explants (mean difference = 0.34, 95% CI [0.03–0.65], p = 0.0239), indicating a partial restoration of the anabolic response (Figure 4d). No effect on transcription of catabolic markers *ADAMTS4* and *ADAMTS4* compared with either Wnt stimulation or α5β1 blockade (Figure 4f,g).

These results suggest that α5β1 integrins can modulate chondrocyte transcription upon Wnt activation and loading. The combined treatment (CHIR99021 + ATN-161 + loading) partially restored load-responsive and anabolic gene expression suppressed by Wnt activation, suggesting that α5β1 integrins contribute to the Wnt-altered mechanoresponse. These findings do not imply that α5β1 blockade is beneficial *per se*, but rather that excessive α5β1 engagement under Wnt⍰activated conditions contributes to maladaptive mechanotransduction.

### Zone-specific actin cytoskeleton reorganization under mechanical load, Wnt activation, and α5β1 integrin blockade

Given that integrins primarily transduce mechanical signals through the cytoskeleton, we next investigated whether the Wnt-induced increase in integrins affects mechanotransduction through F-actin re-organization, and whether α5β1 integrin blockade could modify this response upon Wnt activation. Main effects and a complete set of LMM pairwise comparisons is provided (Supplementary Table 3.5).

In unloaded conditions, treatment did not alter cortical actin levels in either deep (CHIR vs Vehicle: mean difference = 0.0377, 95% CI [−0.0510–0.1264], P = 0.8231; CHIR + ATN vs Vehicle: mean difference = 0.000006, 95% CI [−0.0938–0.0938], P = 1.000) or superficial zone (CHIR vs Vehicle: mean difference = 0.106, 95% CI [0.0198–0.1928], P = 0.066; CHIR + AT vs Vehicle: mean difference = 0.0165, 95% CI [−0.0735–0.1065], P = 0.995), although a trend towards increased cortical actin was observed following Wnt activation in the superficial zone (Figure 5b,l).

**Figure 5.**
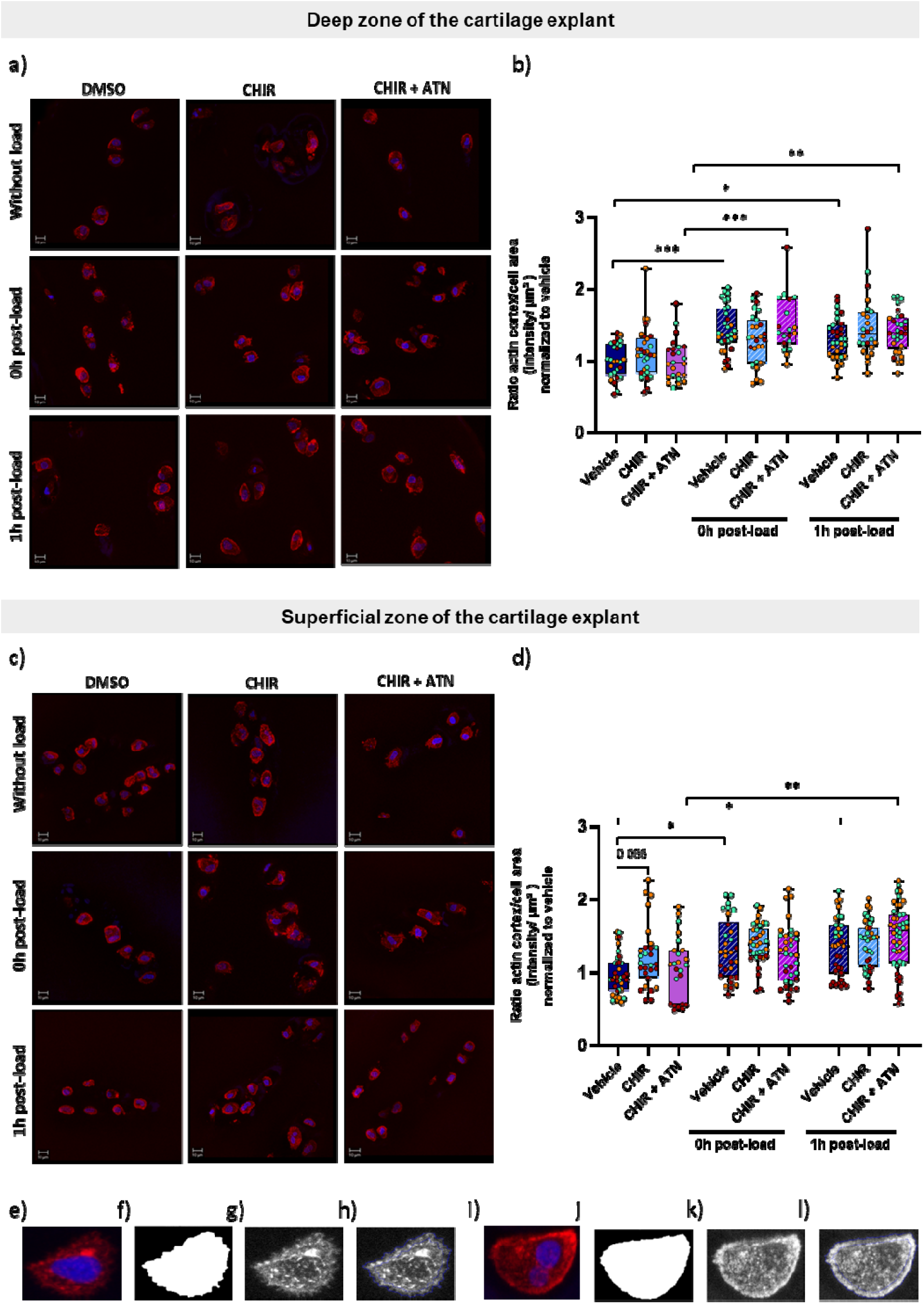
(a,c) Schematic overview of the experimental conditions used to assess cortical actin reorganization in the (a) deep and (c) superficial zones of human articular cartilage explants. Explants were treated with vehicle (DMSO), CHIR99021 (CHIR, 6 μM) to activate canonical Wnt signaling, and/or the α5β1 integrin inhibitor ATN-161 (ATN, 100 μM), under unloaded and mechanically loaded conditions. For each donor, paired explants were either left unloaded or subjected to mechanical loading and collected immediately (0 h post-load) or one hour after stimulation (1 h post-load). (b,d) Quantification of cortical actin organization in the deep and superficial zones across all conditions. (e-h) Representative images of a cell displaying predominantly cytoplasmic actin organization: (e) merged image showing DAPI (blue) and phalloidin (red); (f) cell segmentation used for cell identification; (g) phalloidin channel in greyscale; (h) cortical actin detection as defined by the analysis pipeline. (i–l) Representative images of a cell displaying increased cortical actin organization following mechanical stimulation. Data are presented as individual cells, with colours indicating donor identity and mean ± SEM (n = 3 donors). Statistical analysis was performed using linear mixed-effects models with donor included as a random effect to account for the non-independence of cells derived from the same individual. Cellular measurements were analysed using LMM-effects with donor included as a random effect to account for the non-independence of cells derived from the same individual. Each dot represents an individual cell, and colours indicate donor identity to facilitate visualization of inter-donor variability (n=3).Scale bar: 10 µm. Vehicle = DMSO; CHIR = CHIR99021; ATN = ATN-161; Load = mechanical loading.

However, under loading, distinct changes in F-actin organization were observed across treatments (vehicle, CHIR, and CHIR9 + ATN), both immediately (0h post-load) and 1h post-load. In the deep zone, loading alone induced a more cortical actin distribution compared to the unloaded condition, both at 0h post-load (mean difference = 0.156, 95% CI [0.0648–0.2470], P < 0.0001) and 1h post-load (mean difference = 0.1186, 95% CI [0.0363–0.2009], P = 0.007). However, under Wnt activation, loading did not significantly increase cortical actin at either time point (0h post-load: mean difference = 0.0737, 95% CI [−0.0148–0.1623], P = 0.1617; 1h post-load: mean difference = 0.1340, 95% CI [0.0481–0.2200], = 0.0600) (Figure 5b,l), suggesting that Wnt activation may blunt load-induced cytoskeletal remodelling.

In the **superficial zone**, loading induced time-dependent cytoskeletal reorganization. Immediately after loading (0h post-load), cortical actin distribution was significantly increased (mean difference = 0.0902, 95% CI [0.0043–0.1762], P = 0.0336), whereas neither Wnt activation (CHIR + load vs CHIR; mean difference = 0.0455, 95% CI [−0.0409–0.1318], P = 0.6545) nor combined Wnt activation and α5β1 integrin blockade (mean difference = 0.0553, 95% CI [−0.0385–0.1492], P = 0.5351) had any significant effect. At 1h post-load, cortical actin distribution remained increased compared to unloaded controls (mean difference = 0.1509, 95% CI [0.0737–0.2280], P = 0.0238) with a similar increase observed under combined Wnt activation and α5β1 blockade (mean difference = 0.1261, 95% CI [0.0438–0.2085], P = 0.0020). In contrast, Wnt activation remained non-significant (mean difference = 0.0595, 95% CI [−0.0279–0.1469], P = 0.3704) (Figure 5b,l).

These results indicate that cytoskeletal remodeling is primarily driven by loading but modulated by molecular context. While control conditions show rapid actin reorganization, Wnt activation delays or blunts this response, with α5β1 integrin blockade partially restoring load-induced cytoskeletal adaptation under Wnt activation. These findings suggest that cytoskeletal organisation is an adaptable component of chondrocyte mechanotransduction rather than a sole determinant of mechanoresponsive gene regulation.

## DISCUSSION

Cartilage homeostasis depends on appropriate cellular responses to physiological mechanical forces; understanding how OA-associated signaling pathways disrupt this process is essential for developing therapies to restore tissue dynamics. Studying these interactions in human cartilage provides a unique opportunity to define how the OA environment modifies mechanoresponsiveness. Building on our finding that canonical Wnt activation suppresses load-induced transcription in human cartilage^12^, we investigated underlying mechanisms. Given the critical role of integrins as mechanoreceptors, we first characterised their expression in OA and non-OA cartilage and examined how pharmacological Wnt activation modulates this during loading. *ITGA1, ITGA5* and *ITGB1* were particularly load-responsive. As α5β1 integrin is integral to force transmission and cytoskeletal organisation in chondrocytes, we focused on this receptor to determine whether Wnt activation enhances α5β1 complex formation and whether its blockade can mitigate Wnt-induced alterations in mechanoresponse.

OA chondrocytes exhibited increased *ITGA5* and *ITGB1* but reduced *ITGA10* expression, indicating a disease-associated shift in integrin expression profile consistent with previous reports. OA cartilage expresses integrins rarely detected in healthy tissue, while normally expressed integrins become dysregulated^22^. α5β1 is a key mechanosensor in OA, activated by fibronectin fragments generated during cartilage degradation. Although α5β1 mediates mechanotransduction in both healthy and OA chondrocytes, downstream signaling diverges in OA, activating IL-1β–dependent catabolic pathways. Consistent with this, OA chondrocytes in our study displayed increased α5β1 complex abundance aligning with our previous observation of an amplified acute mechanoresponse in OA cartilage^12^. Increased α5β1availability may contribute to this heightened response, supporting the concept of an early compensatory anabolic phase in OA^23,24^.

Canonical Wnt activation is elevated in OA cartilage and following mechanical injury^17,25^, but its interaction with integrin-mediated adhesion in mechanoresponses remains unclear. Here, Wnt activation increased *ITGA5* and *ITGB1* expression during loading and enhanced α5β1 complex formation, identifying α5β1 as a Wnt-responsive integrin complex in human cartilage. Similar interactions between Wnt signalling and integrin-mediated adhesion have been reported in other mechanosensitive tissues, where Wnt modulates β1-integrin availability and focal adhesion organization^26,27^. In cartilage^26^, Wnt activation also increases matrix stiffness and FAK phosphorylation^28^, supporting its role in modulating integrin–ECM coupling and chondrocyte mechanosensitivity.

Wnt activation dampened the anabolic response to loading in non-OA cartilage, reducing *COL2A1* and *ACAN* expression while increasing *ADAMTS5*, whereas *c-JUN* remained load-responsive. This indicates that chondrocytes still perceive mechanical input but shift toward a catabolic profile. Given the interconnected signaling environment in OA^23^, modulating Wnt alone is unlikely to restore physiological mechanoresponses. Targeting integrin-mediated adhesion provides a mechanistically focused approach to dissect how altered ECM–cell coupling contributes to impaired mechanosensitivity under Wnt activation.

Blocking α5β1 during loading reduced anabolic and mechanoresponsive gene expression whilst increasing catabolic markers, indicating its role in maintaining an anabolic response to physiological loading. Notably, combining α5β1 blockade with Wnt activation partially restored mechanoresponsiveness, with increased *c-JUN* and *COL2A1* expression and a trend towards increased *SOX9*. These findings suggest that excessive α5β1 engagement under Wnt activation disrupts mechanical signaling and that reducing α5β1 availability can partially restore a more typical loading response. The incomplete recovery following α5β1 blockade suggests additional mechanisms contribute to Wnt-induced alterations in mechanoresponsiveness. Compensatory integrin signaling may shift towards others such as αvβ3^29^. Alternatively, canonical Wnt may engage non-canonical pathways, including *WNT5A* and CaMKII-dependent signaling^30–32^, regulating cytoskeletal organisation and adhesion dynamics and may sustain altered mechanoresponses.

As integrins transmit forces to the cytoskeleton and Wnt activation increased α5β1 abundance and altered load-induced transcription, cortical F-actin dynamics provide insight into force transmission. Loading induced a rapid shift toward a more cortical F-actin distribution across all conditions, consistent with rapid, reversible actin remodeling^15,16^ Cell-to-cell variability likely reflects the transient nature of actin reorganisation and cells captured at different stages of remodeling^33^.

Notably, although not significant, Wnt activation showed a trend towards increased cortical actin organization under unloaded conditions, potentially preconditioning the cytoskeleton and contributing to a blunted response to subsequent loading. Importantly, increased cortical actin does not necessarily reflect an effective mechanoresponse, and while required for force transmission, mechanotransduction depends on its capacity for load-induced remodeling. As cortical actin is present in both loaded and unloaded chondrocytes, it likely represents a permissive structural state rather than a direct marker of mechanosensitivity^33–35^. Under Wnt-activation, α5β1 integrin blockade restored load-induced actin organization alongside partial recovery of mechanoresponsive gene expression (*C-JUN* and *COL2A1*), suggesting Wnt primarily affects cytoskeletal adaptability downstream of integrin-mediated adhesion. Load-induced F-actin remodeling was more pronounced and less variable in deep-zone versus superficial-zone chondrocytes, consistent with reported zonal differences in cartilage mechanoresponsiveness^36^.

This study has limitations. α5β1 complex formation was assessed in isolated chondrocytes, preventing *in situ* evaluation of integrin spatial organization. Only short-term mechanoresponses were examined and longer loading may reveal additional mechanisms. Future studies should validate these findings *in situ* and under prolonged loading regimes.

## CONCLUSION

This study identifies α5β1 integrin–mediated adhesion as a key interface through which canonical Wnt signaling modulates chondrocyte mechanoresponsiveness in human cartilage. Wnt activation altered integrin expression and engagement, influencing responses to loading. Although load-induced actin reorganization occurred across conditions, Wnt-associated changes arose primarily from integrin-dependent mechanisms rather than baseline alterations. Together these findings provide mechanistic insight into how dysregulated Wnt-integrin crosstalk may contribute to altered mechanoregulation in osteoarthritis.

## Role of the funding source

This work was supported by Flanders Research Foundation (FWO-Vlaanderen) FWO PhD fellowship (SB/1S18023N) and senior clinical investigator fellowship (18B2122N), KU Leuven/FWO Happy Joints project (C14/24/117 and G045320N) and the EOS excellence program Joint-Against OA (G0F8218N). None of the funding sources had a role in the study design, collection, analysis and interpretation of data; in the writing of the manuscript; and in the decision to submit the manuscript for publication.

## CRediT authorship contribution statement

**N. Viudes-Sarrión:** experimental design; acquisition, analysis and interpretation of data; drafting the manuscript; final approval of the version to be published; agreement to be accountable for all aspects of the work. **R. Castro-Viñuelas:** conception of the work; analysis and interpretation of data; critical reviewing for important intellectual content; final approval of the version to be published; agreement to be accountable for all aspects of the work. **E. Blain:** conception of the work; interpretation of the data; critical reviewing for important intellectual content; final approval of the version to be published; agreement to be accountable for all aspects of the work. **N. Vaes:** analysis of data for the work; final approval of the version to be published; agreement to be accountable for all aspects of the work. **RJ. Lories:** conception of the work; interpretation of the data; critical reviewing for important intellectual content; final approval of the version to be published; agreement to be accountable for all aspects of the work. **I. Jonkers:** conception of the work; interpretation of the data; critical reviewing for important intellectual content; final approval of the version to be published; agreement to be accountable for all aspects of the work.

## Declaration of Competing Interest

None.

## Declaration of use of Generative AI in scientific writing

During the preparation of this work the authors used ChatGPT in order to improve the language. After using this tool, the authors reviewed and edited the content as needed and take full responsibility for the content of the publication.

## Acknowledgments

We are grateful to the UZ Leuven nursing staff for their efforts to provide cartilage samples for this work. We also thank Dr Anthony Hayes and Dr Sophie Gilbert for their valuable assistance with cytoskeletal data acquisition and analysis.

## Notes

### Competing Interest Statement

The authors have declared no competing interest.

