## Supplementary figures and tables for "Interplay between canonical Wnt signaling and α5β1 integrins modulates mechanoresponse in human articular cartilage"

**Supplementary tables**

**Supplementary Table 1. Patients' data**

**1.1 Experiment for integrin expression in OA vs non OA**

| Type | Age (years) | Gender |
|---|---|---|
| Non- OA | 75 | Female |
| Non- OA | 73 | Female |
| Non- OA | 81 | Male |
| OA | 76 | Male |
| OA | 72 | Male |
| OA | 80 | Female |

**1.2 Experiment effects of CHIR on integrin expression**

| Type | Age (years) | Gender |
|---|---|---|
| Non- OA | 82 | Female |
| Non- OA | 85 | Female |
| Non- OA | 77 | Female |
| Non- OA | 97 | Female |
| Non- OA | 75 | Female |
| Non- OA | 60 | Male |
| Non- OA | 85 | Male |

**1.3 Experiment for PLA assay**

| Type | Age (years) | Gender |
|---|---|---|
| Non- OA | 84 | Female |
| Non- OA | 83 | Male |
| Non- OA | 87 | Female |
| OA | 72 | Female |

| | | |
|---|---|---|
| OA | 62 | Female |
| OA | 67 | Male |

**1. 4 Experiment investigating the contribution of integrin α5β1 to Wnt-mediated mechanoresponse**

| Type | Age (years) | Gender |
|---|---|---|
| Non- OA | 82 | Female |
| Non- OA | 87 | Female |
| Non- OA | 83 | Male |
| Non- OA | 87 | Female |
| Non- OA | 82 | Female |
| Non- OA | 68 | Male |
| Non- OA | 80 | Male |

**1.5 Experiment to investigate the contribution of integrin α5β1 and Wnt-mediated cytoskeletal reorganization**

| Type | Age (years) | Gender |
|---|---|---|
| Non- OA | 84 | Male |
| Non- OA | 74 | Female |
| Non- OA | 82 | Female |

**1.6 Experiment for prestoblue and Dose-dependent effects of ATN-161**

| Type | Age (years) | Gender |
|---|---|---|
| Non- OA | 77 | Female |
| Non- OA | 84 | Female |
| Non- OA | 85 | Female |

**Supplementary Table 2. Human primers used in qPCR**

| Primer name | Sequence (5'→ 3') |
|---|---|
| hACAN_fw | ATCCGAGACACCAACGAGAC |
| hACAN_rv | CACTCATTGGCTGCTTCCTG |
| hCFOS_fw | TAGCAAAACGCATGGAGTGT |
| hCFOS_rv | GCCTGGCTCAACATGCTACT |
| hCJUN_fw | AGGGACCCATGGAAGTTTTT |
| hCJUN_rv | TTTTTCTAGGAGTTGTCAATTCAAA |
| hCOL2A1_fw | TGGCAGAGATGGAGAACCTG |
| hCOL2A1_rv | CATCAAATCCTCCAGCCATC |
| hSOX9_fw | TTAAACCCTCTTCAGAGCAAGC |
| hSOX9_rv | GAT TTA GCA CAC TGA TCA CAC G |
| hS29_fw | GGGTCACCAGCAGCTGTACT |
| hS29_rv | AAACACTGGCGGCACATATT |
| hTCF1_fw | CCCCCAACTCTCTCTCTACGA |
| hTCF1_rv | TGCCTGAGGTCAGGGAGTAG |
| hADAMTS4_fw | TGC CCG CTT CAT CAC TGA |
| hADAMTS4_rv | CAA TGG AGC CTC TGG TTT GTC |
| hADAMTS5_fw | TAA GCC CTG GTC CAA ATG C |
| hADAMTS5_rv | AGG TCC AGC AAA CAG TTA CCA |
| hITGB1_fw | GGCAAGTGTGAATGTGGCAG |
| hITGB1_rv | GACTCAATCTCGTCACGGCA |
| hITGB3_fw | GGCAAGTGTGAATGTGGCAG |
| hITGB3_rv | GACTCAATCTCGTCACGGCA |
| hITGA1_fw | CAGCCCCACATTTCAAGTCGT |
| hITGA1_rv | ACCTGTGTCTGTTAGGACCA |
| hITGA2_fw | GCAACTGGTTACTGGTTGGTT |
| hITGA2_rv | GAGGCTCATGTTGGTTTTCATCT |
| hITGA3_fw | TAATACGACTCACTATAGGGAGACCTGCCA |
| hITGA3_rv | TAATACGACTCACTATAGGGAGACGACCCT |
| hITGA4_fw | GCATACAGGTGTCCAGCAGAGA |
| hITGA4_rv | AGGACCAAGGTGGTAAGCAGCT |
| hITGA5_fw | GCCTGTGGAGTACAAGTCCTT |
| hITGA5_rv | AATTCGGGTGAAGTTATCTGTGG |
| hITGA6_fw | CGAAACCAAGGTTCTGAGCCCA |
| hITGA6_rv | CTTGGATCTCCACTGAGGCAGT |
| hITGA10_fw | AATGAGCCTGGAGCCTATCA |
| hITGA10_rv | TATACACGGCTCCTCCGTGA |
| hITGAV_fw | GCTGTCGGAGATTTCAATGGT |
| hITGAV_rv | TCTGCTCGCCAGTAAAATTGT |

**Supplementary Table 3. Statistical analysis details**

**3.1 Experiments integrin expression OA vs non-OA**

| GENE | COMPARISON | ESTIMATE | 95% CI | P VALUE |
|---|---|---|---|---|
| ***ITGB1*** | OA vs non-OA | 0.340 | [0.014 – 0.666] | 0.0445 |
| ***ITGB3*** | OA vs non-OA | 0.036 | [−0.083 – 0.154] | 0.3297 |
| ***ITGA1*** | OA vs non-OA | 0.124 | [−0.374 – 0.621] | 0.4266 |
| ***ITGA2*** | OA vs non-OA | −0.225 | [−0.545 – 0.096] | 0.0946 |
| ***ITGA4*** | OA vs non-OA | −0.001 | [−0.008 – 0.006] | 0.6717 |
| ***ITGA5*** | OA vs non-OA | 0.649 | [0.087 – 1.210] | 0.0327 |
| ***ITGA6*** | OA vs non-OA | −0.026 | [−0.083 – 0.030] | 0.2629 |
| ***ITGA10*** | OA vs non-OA | −0.308 | [−0.510 – −0.106] | 0.0141 |
| ***ITGAV*** | OA vs non-OA | −0.055 | [−0.761 – 0.651] | 0.8266 |

**3.2 Experiment effects of CHIR on integrin expression**

**Main effects**

| GENE | EFFECT | T VALUE | P VALUE |
|---|---|---|---|
| ***TCF1*** | CHIR | 9.675 | 0.0001 |
| | LOAD | 3.325 | 0.0032 |
| | CHIR × LOAD | -2.223 | 0.0373 |
| ***ITGB3*** | CHIR | -0.183 | 0.858 |
| | LOAD | -0.360 | 0.725 |
| | CHIR × LOAD | 1.162 | 0.268 |
| ***ITGB1*** | CHIR | 2.407 | 0.0331 |
| | LOAD | 2.321 | 0.0387 |
| | CHIR × LOAD | 0.547 | 0.0059 |
| ***ITGAV*** | CHIR | 1.452 | 0.172 |
| | LOAD | -1.021 | 0.327 |
| | CHIR × LOAD | 0.061 | 0.952 |
| ***ITGA1*** | CHIR | 2.203 | 0.0478 |
| | LOAD | 3.476 | 0.0045 |
| | CHIR × LOAD | 0.293 | 0.0777 |
| ***ITGA2*** | CHIR | -3.389 | 0.0053 |
| | LOAD | 0.756 | 0.4640 |

| | | | |
|---|---|---|---|
| | CHIR × LOAD | 1.496 | 0.1604 |
| ***ITGA5*** | CHIR | 2.992 | 0.0086 |
| | LOAD | -0.617 | 0.5456 |
| | CHIR × LOAD | 0.665 | 0.5156 |
| ***ITGA6*** | CHIR | -1.560 | 0.145 |
| | LOAD | -1.504 | 0.158 |
| | CHIR × LOAD | 0.759 | 0.463 |
| ***ITGA10*** | CHIR | -0.832 | 0.422 |
| | LOAD | -1.634 | 0.128 |
| | CHIR × LOAD | 0.307 | 0.764 |

**Pairwise comparisons for *TCF-1* (Linear mixed model, emmeans)**

| COMPARISON | ESTIMATE | 95% CI | P VALUE |
|---|---|---|---|
| CHIR vs VEHICLE | +0.7234 | [0.5010 – 0.946] | <0.0001 |
| VEHICLE + LOAD vs VEHICLE | +0.2486 | [0.0261 – 0.471] | 0.0245 |
| CHIR + LOAD vs VEHICLE | +0.7369 | [0.5145 – 0.959] | <0.0001 |
| VEHICLE + LOAD vs CHIR | −0.4748 | [−0.6973 – −0.252] | <0.0001 |
| CHIR + LOAD vs CHIR | +0.0135 | [−0.2090 – 0.236] | 0.9983 |
| CHIR + LOAD vs LOAD | +0.4883 | [0.2658 – 0.711] | <0.0001 |

**Pairwise comparisons for *ITGA1* (Linear mixed model, emmeans)**

| COMPARISON | ESTIMATE | 95% CI | P VALUE |
|---|---|---|---|
| CHIR vs VEHICLE | +0.149 | [−0.005 – 0.373] | 0.0635 |
| VEHICLE + LOAD vs VEHICLE | +0.235 | [0.012 – 0.471] | 0.0374 |
| CHIR + LOAD vs VEHICLE | +0.413 | [0.189 – 0.636] | 0.0004 |
| VEHICLE + LOAD vs CHIR | +0.086 | [−0.138 – 0.310] | 0.6933 |
| CHIR + LOAD vs CHIR | +0.263 | [0.040 – 0.487] | 0.0184 |
| CHIR + LOAD vs LOAD | +0.177 | [−0.046 – 0.401] | 0.1477 |

**Pairwise comparisons for *ITGA2* (Linear mixed model, emmeans)**

| COMPARISON | ESTIMATE | 95% CI | P VALUE |
|---|---|---|---|
| CHIR vs VEHICLE | −0.189 | [−0.373 – −0.005] | 0.0434 |

| COMPARISON | ESTIMATE | 95% CI | P VALUE |
|---|---|---|---|
| VEHICLE + LOAD vs VEHICLE | +0.042 | [−0.142 – 0.226] | 0.9123 |
| CHIR + LOAD vs VEHICLE | −0.029 | [−0.213 – 0.155] | 0.9691 |
| VEHICLE + LOAD vs CHIR | +0.231 | [0.047 – 0.415] | 0.0118 |
| CHIR + LOAD vs CHIR | +0.160 | [−0.024 – 0.344] | 0.1003 |
| CHIR + LOAD vs LOAD | −0.071 | [−0.255 – 0.113] | 0.6930 |

**Pairwise comparisons for *ITGA5* (Linear mixed model, emmeans)**

| COMPARISON | ESTIMATE | 95% CI | P VALUE |
|---|---|---|---|
| CHIR vs VEHICLE | +0.229 | [−0.024 – 0.483] | 0.0583 |
| VEHICLE + LOAD vs VEHICLE | −0.047 | [−0.300 – 0.206] | 0.9493 |
| CHIR + LOAD vs VEHICLE | +0.254 | [0.001 – 0.507] | 0.0491 |
| VEHICLE + LOAD vs CHIR | −0.277 | [−0.530 – −0.023] | 0.0299 |
| CHIR + LOAD vs CHIR | +0.025 | [−0.228 – 0.278] | 0.9921 |
| CHIR + LOAD vs LOAD | +0.301 | [0.048 – 0.555] | 0.0171 |

**Pairwise comparisons for *ITGA6* (Linear mixed model, emmeans)**

| COMPARISON | ESTIMATE | 95% CI | P VALUE |
|---|---|---|---|
| CHIR vs VEHICLE | −0.244 | [−0.760 – 0.272] | 0.5457 |
| VEHICLE + LOAD vs VEHICLE | −0.235 | [−0.751 – 0.281] | 0.5744 |
| CHIR + LOAD vs VEHICLE | −0.311 | [−0.827 – 0.205] | 0.3439 |
| VEHICLE + LOAD vs CHIR | +0.009 | [−0.507 – 0.525] | 1.0000 |
| CHIR + LOAD vs CHIR | −0.067 | [−0.583 – 0.449] | 0.9816 |
| CHIR + LOAD vs LOAD | −0.076 | [−0.592 – 0.440] | 0.9739 |

**Pairwise comparisons for *ITGA10* (Linear mixed model, emmeans)**

| CHIR vs VEHICLE | ESTIMATE | 95% CI | P VALUE |
|---|---|---|---|
| VEHICLE + LOAD vs VEHICLE | −0.098 | [−0.489 – 0.292] | 0.8876 |
| CHIR + LOAD vs VEHICLE | −0.193 | [−0.584 – 0.198] | 0.5085 |
| VEHICLE + LOAD vs CHIR | −0.240 | [−0.631 – 0.150] | 0.3276 |
| CHIR + LOAD vs CHIR | −0.095 | [−0.486 – 0.296] | 0.8976 |

| CHIR vs VEHICLE | ESTIMATE | 95% CI | P VALUE |
|---|---|---|---|
| CHIR + LOAD vs LOAD | −0.142 | [−0.533 – 0.249] | 0.7297 |
| CHIR vs VEHICLE | −0.047 | [−0.438 – 0.344] | 0.9855 |

**Pairwise comparisons for *ITGAV* (Linear mixed model, emmeans)**

| COMPARISON | ESTIMATE | 95% CI | P VALUE |
|---|---|---|---|
| CHIR vs VEHICLE | +0.117 | [−0.149 – 0.384] | 0.5916 |
| VEHICLE + LOAD vs VEHICLE | −0.082 | [−0.349 – 0.184] | 0.8128 |
| CHIR + LOAD vs VEHICLE | +0.042 | [−0.225 – 0.308] | 0.9692 |
| VEHICLE + LOAD vs CHIR | −0.200 | [−0.466 – 0.067] | 0.1823 |
| CHIR + LOAD vs CHIR | −0.075 | [−0.342 – 0.191] | 0.8489 |
| CHIR + LOAD vs LOAD | +0.124 | [−0.142 – 0.391] | 0.5572 |

**Pairwise comparisons for *ITGB3* (Linear mixed model, emmeans)**

| COMPARISON | ESTIMATE | 95% CI | P VALUE |
|---|---|---|---|
| CHIR vs VEHICLE | −0.023 | [−0.430 – 0.385] | 0.9985 |
| VEHICLE + LOAD vs VEHICLE | −0.044 | [−0.452 – 0.363] | 0.9891 |
| CHIR + LOAD vs VEHICLE | +0.136 | [−0.272 – 0.543] | 0.7773 |
| VEHICLE + LOAD vs CHIR | −0.022 | [−0.429 – 0.386] | 0.9987 |
| CHIR + LOAD vs CHIR | +0.158 | [−0.249 – 0.566] | 0.6879 |
| CHIR + LOAD vs LOAD | +0.180 | [−0.227 – 0.588] | 0.5972 |

**Pairwise comparisons for *ITGB1* (Linear mixed model, emmeans)**

| COMPARISON | ESTIMATE | 95% CI | P VALUE |
|---|---|---|---|
| CHIR vs VEHICLE | +0.180 | [−0.428 – 0.670] | 0.0400 |
| VEHICLE + LOAD vs VEHICLE | +0.174 | [−0.074 – 0.421] | 0.2253 |
| CHIR + LOAD vs VEHICLE | +0.412 | [0.165 – 0.659] | 0.0011 |
| VEHICLE + LOAD vs CHIR | −0.006 | [−0.254 – 0.241] | 0.9998 |
| CHIR + LOAD vs CHIR | +0.232 | [0.016 – 0.479] | 0.0404 |
| CHIR + LOAD vs LOAD | +0.238 | [0.010 – 0.486] | 0.0412 |

#### 3.3 Experiment for α5β1 PLA assay

**Main effects**

| EFFECT | T VALUE | P VALUE |
|---|---|---|
| OA x nonOA | 7.749 | 1.03e-12 |
| CHIR x Vehicle | 6.066 | 1.28e-08 |

**Pairwise comparisons for α5β1 PLA assay**

| COMPARISON | ESTIMATE | 95% CI | P VALUE |
|---|---|---|---|
| OA vs NON-OA | +30.1 | [20.36 – 39.84] | 0.0008 |
| CHIR vs VEHICLE | +20.3 | [13.65 – 26.95] | < 0.0001 |

#### 3.4 Experiment investigating the contribution of integrin α5β1 to Wnt-mediated mechanoresponse

**Main effects**

| GENE | EFFECT | T VALUE | P VALUE |
|---|---|---|---|
| ***CJUN*** | CHIR | 2.352 | 0.023435 |
| | ATN | -4.221 | 0.000127 |
| | LOAD | 7.559 | 0.00001 |
| | CHIR × ATN | -0.106 | 0.916045 |
| | CHIR × LOAD | -3.467 | 0.01227 |
| | ATN × LOAD | -4.090 | 0.00191 |
| | CHIR × ATN × LOAD | 4.545 | 0.00001 |
| ***SOX9*** | CHIR | -1.404 | 0.16769 |
| | ATN | -1.674 | 0.10163 |
| | LOAD | 2.761 | 0.00851 |
| | CHIR × ATN | 0.565 | 0.57522 |
| | CHIR × LOAD | -0.277 | 0.78319 |
| | ATN × LOAD | -0.475 | 0.63727 |
| | CHIR × ATN × LOAD | 1.397 | 0.01696 |
| ***COL2*** | CHIR | -1.593 | 0.11859 |
| | ATN | -2.733 | 0.00916 |
| | LOAD | 3.010 | 0.00440 |

| GENE | EFFECT | T VALUE | P VALUE |
|---|---|---|---|
| | CHIR × ATN | 0.612 | 0.54358 |
| | CHIR × LOAD | -3.155 | 0.00297 |
| | ATN × LOAD | -2.096 | 0.04219 |
| | CHIR × ATN × LOAD | 4.301 | 9.93e-05 |
| ***ACAN*** | CHIR | -3.620 | 0.000786 |
| | ATN | -4.042 | 0.000221 |
| | LOAD | 0.619 | 0.539257 |
| | CHIR × ATN | 2.582 | 0.013413 |
| | CHIR × LOAD | -0.086 | 0.931643 |
| | ATN × LOAD | 0.088 | 0.930414 |
| | CHIR × ATN × LOAD | 0.591 | 0.557654 |
| ***ADAMS4*** | CHIR | 3.235 | 0.00237 |
| | ATN | 4.763 | 2.29e-05 |
| | LOAD | 0.071 | 0.94386 |
| | CHIR × ATN | -3.204 | 0.00258 |
| | CHIR × LOAD | -0.128 | 0.89896 |
| | ATN × LOAD | -0.174 | 0.86289 |
| | CHIR × ATN × LOAD | 0.028 | 0.97812 |
| ***ADAMS5*** | CHIR | 3.944 | 0.000299 |
| | ATN | 6.533 | 6.80e-08 |
| | LOAD | 0.610 | 0.544937 |
| | CHIR × ATN | -4.406 | 7.13e-05 |
| | CHIR × LOAD | -0.179 | 0.858906 |
| | ATN × LOAD | -1.631 | 0.110320 |
| | CHIR × ATN × LOAD | 0.637 | 0.527631 |

**Pairwise comparisons for *c-JUN* (Linear mixed model, emmeans)**

| COMPARISON | ESTIMATE | 95% CI | P VALUE |
|---|---|---|---|
| CHIR vs VEHICLE | +0.14 | [−0.07 – 0.35] | 0.4003 |

| COMPARISON | ESTIMATE | 95% CI | P VALUE |
|---|---|---|---|
| ATN vs VEHICLE | −0.26 | [−0.47 – −0.05] | 0.0074 |
| CHIR + ATN vs VEHICLE | −0.12 | [−0.33 – 0.09] | 0.5949 |
| VEHICLE + LOAD vs VEHICLE | +0.46 | [0.25 – 0.67] | <0.0001 |
| CHIR + LOAD vs VEHICLE | +0.31 | [0.09 – 0.52] | 0.0008 |
| ATN + LOAD vs VEHICLE | −0.15 | [−0.36 – 0.06] | 0.3505 |
| CHIR + ATN + LOAD vs VEHICLE | +0.24 | [0.03 – 0.45] | 0.0155 |
| CHIR + LOAD vs VEHICLE + LOAD | −0.16 | [−0.37 – 0.06] | 0.2988 |
| ATN + LOAD vs VEHICLE + LOAD | −0.61 | [−0.82 – −0.40] | <0.0001 |
| CHIR + ATN + LOAD vs VEHICLE + LOAD | −0.22 | [−0.43 – −0.01] | 0.0348 |
| ATN vs CHIR | −0.40 | [−0.61 – −0.19] | <0.0001 |
| CHIR + ATN vs CHIR | −0.27 | [−0.48 – −0.06] | 0.0049 |
| VEHICLE + LOAD vs CHIR | −0.32 | [−0.53 – −0.11] | 0.0004 |
| CHIR + LOAD vs CHIR | +0.16 | [−0.05 – 0.37] | 0.2522 |
| ATN + LOAD vs CHIR + LOAD | −0.45 | [−0.67 – −0.24] | <0.0001 |
| CHIR + ATN + LOAD vs CHIR + LOAD | −0.07 | [−0.28 – 0.15] | 0.9763 |
| CHIR + ATN vs ATN | +0.13 | [−0.08 – 0.35] | 0.4855 |
| VEHICLE + LOAD vs ATN | +0.72 | [0.51 – 0.93] | <0.0001 |
| ATN + LOAD vs ATN | +0.11 | [−0.10 – 0.32] | 0.7361 |
| CHIR + ATN + LOAD vs ATN + LOAD | +0.39 | [0.18 – 0.60] | <0.0001 |
| VEHICLE + LOAD vs CHIR + ATN | +0.58 | [0.37 – 0.79] | <0.0001 |
| CHIR + ATN + LOAD vs CHIR + ATN | +0.36 | [0.15 – 0.57] | <0.0001 |

**Pairwise comparisons for *SOX9* (Linear mixed model, emmeans)**

| COMPARISON | ESTIMATE | 95% CI | P VALUE |
|---|---|---|---|
| CHIR vs VEHICLE | −0.17 | [−0.58 – 0.25] | 0.9014 |
| ATN vs VEHICLE | −0.20 | [−0.61 – 0.21] | 0.7889 |
| CHIR + ATN vs VEHICLE | −0.27 | [−0.68 – 0.14] | 0.4411 |
| VEHICLE + LOAD vs VEHICLE | +0.33 | [−0.08 – 0.74] | 0.0606 |
| CHIR + LOAD vs VEHICLE | +0.12 | [−0.30 – 0.53] | 0.9865 |
| ATN + LOAD vs VEHICLE | +0.05 | [−0.36 – 0.46] | 0.9999 |

| COMPARISON | ESTIMATE | 95% CI | P VALUE |
|---|---|---|---|
| CHIR + ATN + LOAD vs VEHICLE | +0.26 | [−0.15 – 0.68] | 0.4787 |
| CHIR + LOAD vs VEHICLE + LOAD | −0.21 | [−0.63 – 0.20] | 0.7247 |
| ATN + LOAD vs VEHICLE + LOAD | −0.28 | [−0.69 – 0.13] | 0.4039 |
| CHIR + ATN + LOAD vs VEHICLE + LOAD | −0.07 | [−0.48 – 0.35] | 0.9996 |
| ATN vs CHIR | −0.03 | [−0.45 – 0.38] | 1.0000 |
| CHIR + ATN vs CHIR | −0.10 | [−0.52 – 0.31] | 0.9924 |
| VEHICLE + LOAD vs CHIR | +0.50 | [0.08 – 0.91] | 0.0086 |
| CHIR + LOAD vs CHIR | +0.28 | [−0.13 – 0.70] | 0.3908 |
| ATN + LOAD vs CHIR + LOAD | −0.07 | [−0.48 – 0.35] | 0.9996 |
| CHIR + ATN + LOAD vs CHIR + LOAD | +0.15 | [−0.26 – 0.56] | 0.9447 |
| CHIR + ATN vs ATN | −0.07 | [−0.49 – 0.34] | 0.9993 |
| VEHICLE + LOAD vs ATN | +0.53 | [0.12 – 0.94] | 0.0041 |
| ATN + LOAD vs ATN | +0.25 | [−0.16 – 0.66] | 0.5525 |
| CHIR + ATN + LOAD vs ATN + LOAD | +0.21 | [−0.20 – 0.05] | 0.0623 |
| VEHICLE + LOAD vs CHIR + ATN | +0.60 | [0.19 – 1.01] | 0.0007 |

**Pairwise comparisons for *COL2A1* (Linear mixed model, emmeans)**

| COMPARISON | ESTIMATE | 95% CI | P VALUE |
|---|---|---|---|
| CHIR vs VEHICLE | −0.14 | [−0.45 – 0.17] | 0.8271 |
| ATN vs VEHICLE | −0.24 | [−0.55 – 0.07] | 0.2214 |
| CHIR + ATN vs VEHICLE | −0.31 | [−0.62 – 0.00] | 0.0700 |
| VEHICLE + LOAD vs VEHICLE | +0.27 | [−0.04 – 0.58] | 0.0132 |
| CHIR + LOAD vs VEHICLE | −0.27 | [−0.58 – 0.04] | 0.1235 |
| ATN + LOAD vs VEHICLE | −0.24 | [−0.55 – 0.07] | 0.2399 |
| CHIR + ATN + LOAD vs VEHICLE | +0.06 | [−0.24 – 0.37] | 0.9976 |
| CHIR + LOAD vs VEHICLE + LOAD | −0.54 | [−0.85 – −0.23] | <0.0001 |
| ATN + LOAD vs VEHICLE + LOAD | −0.51 | [−0.82 – −0.20] | 0.0001 |
| CHIR + ATN + LOAD vs VEHICLE + LOAD | −0.20 | [−0.51 – 0.11] | 0.4387 |
| ATN vs CHIR | −0.10 | [−0.41 – 0.21] | 0.9658 |
| CHIR + ATN vs CHIR | −0.17 | [−0.48 – 0.14] | 0.6847 |

| COMPARISON | ESTIMATE | 95% CI | P VALUE |
|---|---|---|---|
| VEHICLE + LOAD vs CHIR | +0.41 | [0.10 – 0.72] | 0.0026 |
| CHIR + LOAD vs CHIR | −0.13 | [−0.44 – 0.18] | 0.8851 |
| ATN + LOAD vs CHIR + LOAD | −0.27 | [−0.60 – 0.06] | 0.1871 |
| CHIR + ATN + LOAD vs CHIR + LOAD | +0.34 | [0.03 – 0.65] | 0.0239 |
| CHIR + ATN vs ATN | −0.07 | [−0.34 – 0.24] | 1.0000 |
| VEHICLE + LOAD vs ATN | +0.51 | [0.20 – 0.82] | 0.0001 |
| ATN + LOAD vs ATN | +0.00 | [−0.31 – 0.31] | 1.0000 |
| CHIR + ATN + LOAD vs ATN + LOAD | +0.31 | [−0.01 – 0.61] | 0.0404 |
| VEHICLE + LOAD vs CHIR + ATN | +0.58 | [0.27 – 0.89] | <0.0001 |

**Pairwise comparisons for *ACAN* (Linear mixed model, emmeans)**

| COMPARISON | ESTIMATE | 95% CI | P VALUE |
|---|---|---|---|
| CHIR vs VEHICLE | −0.35 | [−0.68 – −0.02] | 0.0346 |
| ATN vs VEHICLE | −0.39 | [−0.72 – −0.06] | 0.0119 |
| CHIR + ATN vs VEHICLE | −0.39 | [−0.72 – −0.05] | 0.0129 |
| VEHICLE + LOAD vs VEHICLE | +0.06 | [−0.27 – 0.39] | 0.9991 |
| CHIR + LOAD vs VEHICLE | −0.30 | [−0.63 – 0.03] | 0.1052 |
| ATN + LOAD vs VEHICLE | −0.32 | [−0.65 – 0.02] | 0.0724 |
| CHIR + ATN + LOAD vs VEHICLE | −0.21 | [−0.55 – 0.12] | 0.4820 |
| CHIR + LOAD vs VEHICLE + LOAD | −0.41 | [−0.74 – −0.07] | 0.0071 |
| ATN + LOAD vs VEHICLE + LOAD | −0.45 | [−0.78 – −0.11] | 0.0022 |
| CHIR + ATN + LOAD vs VEHICLE + LOAD | −0.17 | [−0.51 – 0.16] | 0.7032 |
| ATN vs CHIR | −0.04 | [−0.37 – 0.29] | 0.9999 |
| CHIR + ATN vs CHIR | −0.04 | [−0.37 – 0.30] | 1.0000 |
| VEHICLE + LOAD vs CHIR | +0.41 | [0.07 – 0.74] | 0.0071 |
| CHIR + LOAD vs CHIR | +0.05 | [−0.29 – 0.38] | 0.9998 |
| ATN + LOAD vs CHIR + LOAD | +0.09 | [−0.25 – 0.42] | 0.9917 |
| CHIR + ATN + LOAD vs CHIR + LOAD | +0.18 | [−0.16 – 0.51] | 0.7205 |
| CHIR + ATN vs ATN | +0.00 | [−0.33 – 0.33] | 1.0000 |
| VEHICLE + LOAD vs ATN | +0.45 | [0.11 – 0.78] | 0.0022 |

| COMPARISON | ESTIMATE | 95% CI | P VALUE |
|---|---|---|---|
| ATN + LOAD vs ATN | +0.07 | [−0.26 – 0.40] | 0.9972 |
| CHIR + ATN + LOAD vs ATN + LOAD | +0.10 | [−0.23 – 0.44] | 0.9979 |
| VEHICLE + LOAD vs CHIR + ATN | +0.44 | [0.11 – 0.78] | 0.0024 |
| CHIR + ATN + LOAD vs CHIR + ATN | +0.17 | [−0.16 – 0.51] | 0.7205 |

**Pairwise comparisons for *ADAMTS4* (Linear mixed model, emmeans)**

| COMPARISON | ESTIMATE | 95% CI | P VALUE |
|---|---|---|---|
| CHIR vs VEHICLE | +0.26 | [−0.02 – 0.53] | 0.0831 |
| ATN vs VEHICLE | +0.38 | [0.10 – 0.65] | 0.0016 |
| CHIR + ATN vs VEHICLE | +0.27 | [0.00 – 0.55] | 0.0496 |
| VEHICLE + LOAD vs VEHICLE | +0.00 | [−0.27 – 0.28] | 1.0000 |
| CHIR + LOAD vs VEHICLE | +0.25 | [−0.03 – 0.52] | 0.1048 |
| ATN + LOAD vs VEHICLE | +0.36 | [0.09 – 0.63] | 0.0027 |
| CHIR + ATN + LOAD vs VEHICLE | +0.25 | [−0.02 – 0.52] | 0.0962 |
| CHIR + LOAD vs VEHICLE + LOAD | −0.25 | [−0.52 – 0.02] | 0.0966 |
| ATN + LOAD vs VEHICLE + LOAD | −0.37 | [−0.64 – −0.10] | 0.0020 |
| CHIR + ATN + LOAD vs VEHICLE + LOAD | −0.27 | [−0.54 – 0.01] | 0.0583 |
| ATN vs CHIR | +0.12 | [−0.15 – 0.39] | 0.8552 |
| CHIR + ATN vs CHIR | +0.02 | [−0.25 – 0.29] | 1.0000 |
| VEHICLE + LOAD vs CHIR | −0.25 | [−0.52 – 0.02] | 0.0966 |
| CHIR + LOAD vs CHIR | −0.01 | [−0.28 – 0.26] | 1.0000 |
| ATN + LOAD vs CHIR + LOAD | +0.12 | [−0.17 – 0.39] | 0.9172 |
| CHIR + ATN + LOAD vs CHIR + LOAD | −0.01 | [−0.28 – 0.27] | 1.0000 |
| CHIR + ATN vs ATN | −0.10 | [−0.38 – 0.17] | 0.9329 |
| VEHICLE + LOAD vs ATN | −0.37 | [−0.64 – −0.10] | 0.0020 |
| ATN + LOAD vs ATN | −0.01 | [−0.29 – 0.26] | 1.0000 |
| CHIR + ATN + LOAD vs ATN + LOAD | −0.11 | [−0.39 – 0.16] | 0.8253 |
| VEHICLE + LOAD vs CHIR + ATN | −0.27 | [−0.54 – 0.01] | 0.0583 |
| CHIR + ATN + LOAD vs CHIR + ATN | −0.02 | [−0.30 – 0.25] | 1.0000 |

**Pairwise comparisons for *ADAMTS5* (Linear mixed model, emmeans)**

| COMPARISON | ESTIMATE | 95% CI | P VALUE |
|---|---|---|---|
| CHIR vs VEHICLE | +0.30 | [0.04 – 0.57] | 0.0154 |
| ATN vs VEHICLE | +0.50 | [0.24 – 0.77] | <0.0001 |
| CHIR + ATN vs VEHICLE | +0.33 | [0.06 – 0.59] | 0.0069 |
| VEHICLE + LOAD vs VEHICLE | +0.05 | [−0.22 – 0.31] | 0.9992 |
| CHIR + LOAD vs VEHICLE | +0.33 | [0.06 – 0.60] | 0.0060 |
| ATN + LOAD vs VEHICLE | +0.37 | [0.11 – 0.64] | 0.0013 |
| CHIR + ATN + LOAD vs VEHICLE | +0.27 | [0.01 – 0.54] | 0.0390 |
| CHIR + LOAD vs VEHICLE + LOAD | −0.26 | [−0.52 – −0.01] | 0.0471 |
| ATN + LOAD vs VEHICLE + LOAD | −0.33 | [−0.72 – −0.19] | 0.0010 |
| CHIR + ATN + LOAD vs VEHICLE + LOAD | −0.28 | [−0.55 – −0.01] | 0.0334 |
| ATN vs CHIR | +0.20 | [−0.07 – 0.47] | 0.2814 |
| CHIR + ATN vs CHIR | +0.02 | [−0.24 – 0.29] | 1.0000 |
| VEHICLE + LOAD vs CHIR | −0.26 | [−0.52 – 0.01] | 0.0671 |
| CHIR + LOAD vs CHIR | +0.03 | [−0.24 – 0.29] | 1.0000 |
| ATN + LOAD vs CHIR + LOAD | +0.04 | [−0.23 – 0.31] | 0.9997 |
| CHIR + ATN + LOAD vs CHIR + LOAD | −0.06 | [−0.32 – 0.21] | 0.9975 |
| CHIR + ATN vs ATN | −0.18 | [−0.44 – 0.09] | 0.4362 |
| VEHICLE + LOAD vs ATN | −0.46 | [−0.72 – −0.19] | <0.0001 |
| ATN + LOAD vs ATN | −0.13 | [−0.40 – 0.14] | 0.7774 |
| CHIR + ATN + LOAD vs ATN + LOAD | −0.10 | [−0.36 – 0.17] | 0.9404 |
| VEHICLE + LOAD vs CHIR + ATN | −0.28 | [−0.55 – −0.01] | 0.0334 |
| CHIR + ATN + LOAD vs CHIR + ATN | −0.05 | [−0.32 – 0.21] | 0.9985 |

### 3.5 Experiment to investigate the contribution of integrin α5β1 and Wnt-mediated cytoskeletal reorganization

#### Main effects

#### DEEP ZONE — JUST AFTER LOADING

| EFFECT | T VALUE | P VALUE |
|---|---|---|
| CHIR | 1.251 | 0.2129 |
| CHIR+ ATN | -1.212 | 0.2272 |

| EFFECT | T VALUE | P VALUE |
|---|---|---|
| LOAD | 5.159 | 7.44e-07 |
| CHIR × LOAD | -1.924 | 0.0562 |
| CHIR+ ATN × LOAD | 2.116 | 0.0360 |

**DEEP ZONE — 1H AFTER LOADING**

| EFFECT | T VALUE | P VALUE |
|---|---|---|
| CHIR | 1.230 | 0.2200 |
| CHIR+ ATN | -1.263 | 0.2080 |
| LOAD | 4.226 | 3.94e-05 |
| CHIR × LOAD | 0.379 | 0.7050 |
| CHIR+ ATN × LOAD | 0.274 | 0.7840 |

**SUPERFICIAL ZONE — JUST AFTER LOADING**

| EFFECT | T VALUE | P VALUE |
|---|---|---|
| CHIR | 3.590 | 0.0004 |
| CHIR+ ATN | -2.752 | 0.0065 |
| LOAD | 3.068 | 0.0025 |
| CHIR × LOAD | -1.089 | 0.2776 |
| CHIR+ ATN × LOAD | 0.227 | 0.8210 |

**SUPERFICIAL ZONE — 1H AFTER LOADING**

| EFFECT | T VALUE | P VALUE |
|---|---|---|
| CHIR | 3.947 | 0.0001 |
| CHIR+ ATN | -3.034 | 0.0027 |
| LOAD | 5.703 | 4.34e-08 |
| CHIR × LOAD | -2.275 | 0.0240 |
| CHIR+ ATN × LOAD | 1.632 | 0.1043 |

**Pairwise comparisons for Deep zone of the cartilage**

| COMPARISON | ESTIMATE | 95% CI | P VALUE |
|---|---|---|---|
| CHIR vs VEHICLE | +0.0377 | [−0.0510 – 0.1264] | 0.8231 |
| CHIR + ATN vs VEHICLE | +0.000006 | [−0.0938 – 0.0938] | 1.0000 |
| CHIR + ATN vs CHIR | −0.0377 | [−0.1290 – 0.0535] | 0.8400 |
| VEHICLE 0h POST-LOAD vs VEHICLE | +0.156 | [0.0648 – 0.2470] | <0.0001 |
| CHIR 0h POST-LOAD vs VEHICLE | +0.111 | [0.0207 – 0.2022] | 0.0380 |
| CHIR + ATN 0h POST-LOAD vs VEHICLE | +0.171 | [0.0693 – 0.2737] | <0.0001 |
| CHIR 0h POST-LOAD vs CHIR | +0.0737 | [−0.0148 – 0.1623] | 0.1617 |
| CHIR + ATN 0h POST-LOAD vs CHIR + ATN | +0.171 | [0.0682 – 0.2747] | 0.0001 |
| VEHICLE 1h POST-LOAD vs VEHICLE | +0.1186 | [0.0363 – 0.2009] | 0.0070 |
| CHIR 1h POST-LOAD vs VEHICLE | +0.1688 | [0.0811 – 0.2564] | 0.0220 |
| CHIR + ATN 1h POST-LOAD vs VEHICLE | +0.1433 | [0.0598 – 0.2268] | 0.0352 |
| CHIR 1h POST-LOAD vs CHIR | +0.1340 | [0.0481 – 0.2200] | 0.0600 |
| CHIR + ATN 1h POST-LOAD vs CHIR + ATN | +0.1454 | [0.0599 – 0.2308] | <0.0001 |

**Pairwise comparisons for Superficial zone of the cartilage**

| COMPARISON | ESTIMATE | 95% CI | P VALUE |
|---|---|---|---|
| CHIR vs VEHICLE | +0.1063 | [0.0198 – 0.1928] | 0.0660 |
| CHIR + ATN vs VEHICLE | +0.0165 | [−0.0735 – 0.1065] | 0.9950 |
| CHIR + ATN vs CHIR | −0.0898 | [−0.1851 – 0.0055] | 0.0772 |
| VEHICLE 0h POST-LOAD vs VEHICLE | +0.0902 | [0.0043 – 0.1762] | 0.0336 |
| CHIR 0h POST-LOAD vs VEHICLE | +0.1518 | [0.0720 – 0.2316] | 0.0100 |
| CHIR + ATN 0h POST-LOAD vs VEHICLE | +0.0718 | [−0.0129 – 0.1565] | 0.1477 |
| CHIR 0h POST-LOAD vs CHIR | +0.0455 | [−0.0409 – 0.1318] | 0.6545 |
| CHIR + ATN 0h POST-LOAD vs CHIR + ATN | +0.0553 | [−0.0385 – 0.1492] | 0.5351 |
| VEHICLE 1h POST-LOAD vs VEHICLE | +0.1509 | [0.0737 – 0.2280] | 0.0238 |
| CHIR 1h POST-LOAD vs VEHICLE | +0.1706 | [0.0888 – 0.2524] | 0.0211 |
| CHIR + ATN 1h POST-LOAD vs VEHICLE | +0.1431 | [0.0702 – 0.2159] | 0.0321 |
| CHIR 1h POST-LOAD vs CHIR | +0.0595 | [−0.0279 – 0.1469] | 0.3704 |
| CHIR + ATN 1h POST-LOAD vs CHIR + ATN | +0.1261 | [0.0438 – 0.2085] | 0.0020 |

**Supplementary figures.**

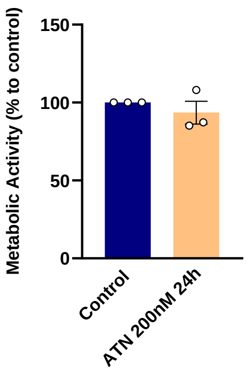


**Figure S.1.** Cartilage explant viability. Each individual data point represents the mean of three technical replicates. Data are presented as individual data points for each donor sample, along with the mean ± SEM (n = 3 per group).

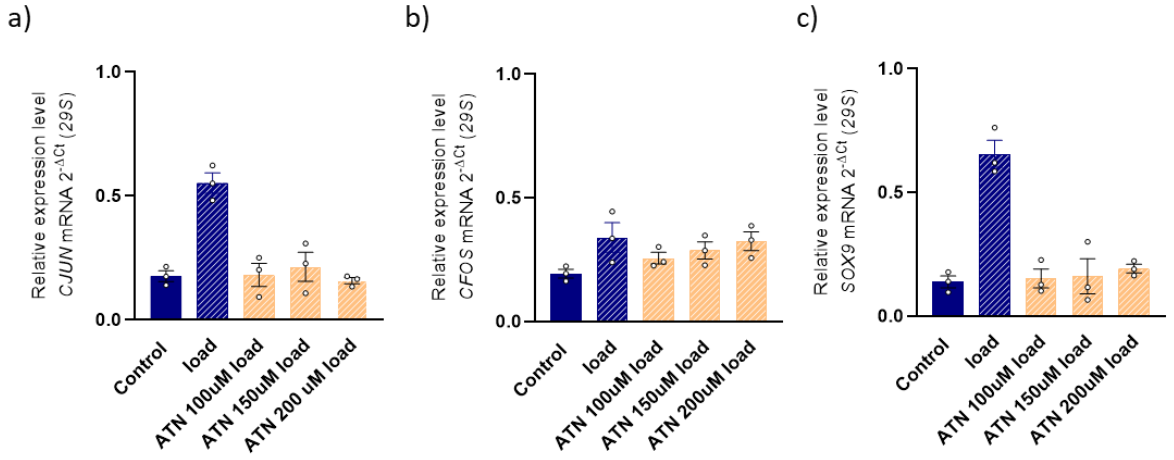


**Figure S.2.** Dose–response analysis of the integrin α5β1 blocker ATN-161 (100 µM, 150 µM, and 200 µM) in mechanically loaded human articular cartilage explants. Gene expression of the mechanoresponsive markers **a)** *cJUN* and **b)** *cFOS*, and the **c)** chondrogenic marker *SOX9*, was assessed following loading to determine the concentration at which ATN-161 begins to alter the mechanoresponsive profile. Data are expressed as individual data points representing each donor and mean ± SEM (n = 3). ATN = ATN-161; Load = mechanical loading.
